# Establishing immortalized brown and white preadipocyte cell lines from young and aged mice

**DOI:** 10.1101/2024.10.10.617572

**Authors:** Xiangdong Wu, Salaheldeen Elsaid, Florian Levet, Winson Li, Sui Seng Tee

## Abstract

Studying adipogenesis and adipocyte biology requires the isolation of primary preadipocytes from adipose tissues. However, primary preadipocytes have a limited lifespan, can only undergo a finite number of divisions, and often lose their original biological characteristics before becoming senescent. The repeated isolation of fresh preadipocytes, particularly from young pups or aged animals, is costly and time-consuming. Immortalization of these cells offers a solution by overcoming cellular senescence and maintaining proliferative capacity, allowing for long-term studies without the continuous need to isolate new cells from animals. Immortalized cell lines thus provide a consistent and reproducible experimental model, significantly reducing variability across different animals. However, successfully establishing immortalized preadipocyte cell lines presents challenges, including selecting appropriate adipose tissue depots, isolating primary preadipocytes, and choosing an effective immortalization strategy. In this study, we present optimized protocols and share first-hand experiences establishing immortalized brown and white preadipocyte cell lines from young and aging mice. These protocols offer a valuable resource for researchers studying adipogenesis, metabolism, and adipocyte biology.

**Support Protocol 1**: Retrovirus production

**Basic Protocol 1**: Isolation and culture of primary brown and white preadipocytes from mouse interscapular brown adipose tissue (iBAT) and subcutaneous white adipose tissue (sWAT) in the same region

**Basic Protocol 2**: Immortalization of mouse brown and white preadipocytes

**Basic Protocol 3**: Selection of immortalized preadipocytes

**Basic Protocol 4**: Selection of single-cell clones of immortalized preadipocytes

**Support Protocol 2**: Cryopreservation of immortalized preadipocytes

**Support Protocol 3**: Wake up and culture of immortalized preadipocytes

**Support Protocol 4**: Subculture and expansion of immortalized preadipocytes

**Basic Protocol 5**: Differentiation of immortalized mouse brown and white preadipocytes

**Support Protocol 5**: Lipid droplet staining and nucleus counterstaining

**Support Protocol 6**: Mitochondria staining and nucleus counterstaining

## INTRODUCTION

Adipocytes play a critical role in energy metabolism. White adipocytes store energy in a single large lipid droplet, while brown adipocytes contain multiple small lipid droplets and a high number of mitochondria, contributing to non-shivering thermogenesis, which is crucial for maintaining body temperature. Due to its thermogenic properties in response to cold and overnutrition, as well as its ability to convert energy into heat, activating brown adipose tissue— unexpectedly discovered in adult humans through radiological approaches—has become an attractive strategy for maintaining energy homeostasis and treating metabolic disorders (Cero et al., 2021).

Beyond their roles in metabolic processes, adipocytes secrete many cytokines, also known as adipokines. Both white and brown adipocytes secrete common cytokines such as adiponectin and IL-6, but they also secrete distinct sets of cytokines. For example, white adipocytes secrete leptin, TNF-α, PAI-1, MCP-1, and angiotensinogen, while brown adipocytes secrete norepinephrine, Nrg4, and VEGF. These adipokines not only play roles in numerous cellular signaling pathways and regulate important metabolic processes, but also trigger immune responses, contributing to inflammation and immune regulation. Adipose tissues are considered active endocrine and immune tissues (Machdo et al., 2022; Clemente-Suarez et al., 2023; Ghesmati et al., 2024). Both white and brown adipocytes are promising therapeutic targets for metabolic disorders such as obesity, type 2 diabetes, cardiovascular diseases, fatty liver, and certain cancers, including breast and pancreatic cancer.

Brown and white adipocytes also play significant roles in aging. As individuals age, the function and thermogenic capacity of brown adipocytes decline, slowing metabolic rates and increasing the risk of insulin resistance, obesity, and metabolic disorders such as type 2 diabetes. Meanwhile, white adipocytes persistently secrete pro-inflammatory cytokines such as TNF-α, IL-6, and MCP-1, leading to a state of chronic low-grade inflammation that contributes to various age-related diseases, including cardiovascular diseases and certain cancers (Rogers et al., 2012; Goncalves et al., 2017; Duteil et al., 2017; Nanduri et al., 2021).

To better understand and investigate the roles of brown and white adipocytes in energy homeostasis, adipogenesis, thermogenesis, metabolic disorders, and aging processes in vitro, we established immortalized mouse brown and white preadipocyte cell lines from young (one-day-old pups) and old mice (12-month-old and 27-month-old). One group of old mice (12 months) was also maintained on a normal diet and a high-fat, high-fructose diet for 10 months.

## RETROVIRUS PRODUCTION

Before isolating a brown or white preadipocyte, the retrovirus that delivers the Large T antigen or Telomerase reverse transcriptase for immortalization must be prepared and in the status of ready to use.

### Materials

HEK-293T cells
pBABE-neo large T cDNA, addgene plasmid #: 1780
pCL-Eco, addgene plasmid #: 12371
pVSV-G, addgene plasmid #: 138479
Opti MEM reduced serum medium (Opti MEM)
Fetal Bovine Serum (FBS)
Retrovirus packing medium: Opti MEM reduced serum medium (with glutamine, pyruvate) with 5% FBS
45μm pore size filter

Day 0: preparation of 293T cells as producer cells

1. Seed 293T cells to be 95-99% confluent at the transfection, approximately 1.2×10^6^ cell/well of a 6-well plate in 2ml packing medium, and transfer the plate back to an incubator with normal cell culture condition (37°C, 5% CO_2_) for overnight (16-24 hours).

Day 1: transfection

2. Dilute Lipo3000 in Opti MEM. mix 250μl Opti MEM with 7μl Lipo3000. (tube A)
3. Dilute retroviral packaging vectors and expression vectors in Opti MEM, then add P3000 enhancer reagent. mix 250μl Opti MEM with 2.25μg packing vector and 0.75μg expression vector and 6μl P3000. (tube B)
4. Add the content of tube A to tube B (1:1 ratio) and mix well.
5. Incubate for 10-20 minutes at room temperature to form a DNA-lipid complex.
6. Remove 50% volume of media (1ml) from each well, now 1ml remaining packing medium in each well.
7. Add DNA-lipid complexes from step 4 and step 5 to cells in each well.
8. Incubate cells for 6 hours at 37°C and 5% CO_2_.
9. Remove medium including DNA-lipid complexes and replace with 2ml fresh packing medium.

Day 2: the first collection of viral production

10. Harvest entire volume of cell supernatant (2ml) at 24 hours post-transfection; store at 4°C.
11. Replace with 2ml fresh pre-warmed packaging medium for each well.
12. Incubate cells overnight at 37 °C and 5% CO_2_. Day 3: the second collection of viral production
13. Harvest entire volume of cell supernatant (2ml) at 52 hours post-transfection; store at 4°C.
14. Centrifuge viral supernatant at 2000 rpm for 5 minutes. The first and second collections of viral production can be combined or kept separate.
15. Filter the supernatant through a 45μm pore size filter.
16. Titer, aliquot and store virus at −80°C.

### Retrovirus Quantitation

We used a commercially available QuickTiter^TM^ retrovirus quantitation kit for viral particle quantification. Protocols can be referenced from manufacturer product manuals.

## ISOLATION AND CULTURE OF PRIMARY BROWN OR WHITE PREADIPOCYTES FROM MOUSE INTERSCAPULAR BROWN ADIPOSE TISSUE (iBAT) or SUBCUTANEOUS WHITE ADIPOSE TISSUE (sWAT) IN THE SAME INTERSCAPULAR REGION

### Lab Animal

Newborn mouse pup (1–2 days old)
20 weeks young adult
12 months aged adult
27 months aged adult

## Lab equipment, Tools, and cell culture supplies

Cell culture incubator
water bath with shaker
Scissors and forceps (autoclaved)
50-ml and 15-ml Falcon tubes
100-μm cell strainers
220-μm syringe-driven filters
50-ml syringes
Cell culture plates (6–12 well)

### Buffer and Culture Medium

1. Pre-digestion buffer

1) Prepare 1 lite of pre-digestion buffer including 123.0 mM NaCl, 5.0 mM KCl, 1.3 mM CaCl2, 5.0mM Glucose, 100.0mM N-2-hydroxyethylpiperazine’-2-ethanesulfonic acid (HEPES).
2) adjust pH to 7.5, and store at 4 °C.
2. Digestion buffer

1) Add 2g of BSA, 50mg of collagenase II, 1ml of promcine to 49ml of pre-digestion buffer.
2) Vortex, make sure BSA and collagenase II are completely dissolved in buffer.
3) Filter the buffer through 220μm filters.
3) Keep the buffer in a 37 °C water bath. Note: The digestion buffer contains 4% (g/v) of BSA, 1mg/ml of collagenase II, and 100μg/ml primocin.
3. Primary preadipocytes growth medium

1) Add 50ml of FBS and 1ml of primocin to 449ml of Dulbecco’s Modified Eagle Medium/F12 (DMEM/F12).
2) Filter the medium through 220μm filters. Note: DMEM/F-12 medium containing 10% FBS and 100μg/ml primocin.
4. Induction medium for brown adipocytes

The induction medium for brown adipocytes containing primary preadipocyte growth medium with 20nM insulin, 0.5mM IBMX, 125nM indomethacin, 1nM T3, 1μM Dexamethasone, and 1μM Rosiglitazone.
1) prepare 0.348mM of insulin stock solution in 0.01N HCl, 50mM IBMX stock solution in 1M KOH, 1.25mM Indomethacin stock solution in ethanol, 29.71μM T3 stock solution in NaOH, 5mM Dexamethasone stock solution in ethanol, 27.97 Rosiglitazone stock solution in DMSO.
2) Add 5.8μl insulin stock, 1ml IBMX stock, 10μl indomethacin stock, 3.4μl T3 stock, 20μl Dexamethasone stock, 3.6 Rosiglitazone stock into 99ml of primary brown preadipocytes growth medium.
3) Filter the medium through 220μm filters.
5. Induction medium for white adipocyte

The induction medium for white adipocytes contains primary preadipocyte growth medium with 20nM insulin, 0.5mM IBMX, 125nM indomethacin, 1μM Dexamethasone, and 1μM Rosiglitazone.
1) Add 5.8μl insulin stock, 1ml IBMX stock, 10μl indomethacin stock, 20μl Dexamethasone stock, 3.6 Rosiglitazone stock into 99ml of primary preadipocytes growth medium.
2) Filter the medium through 220μm filters.
6. Maintenance (or differentiation) medium for brown adipocyte

The maintenance or differentiation medium for brown adipocytes containing primary brown preadipocyte growth medium with 20nM insulin, and 1nM T3.
1) Add 5.8μl insulin stock, 3.4μl T3 stock to 100ml of primary brown preadipocytes growth medium.
2) Filter through 220μm filters.
7. Maintenance (or differentiation) medium for white adipocyte

The maintenance medium for white adipocytes contains primary preadipocyte growth medium with 20nM insulin.
1) Add 5.8μl insulin stock to 100ml of primary brown preadipocyte growth medium.
2) Filter through 220μm filters.
8. Cell freezes medium

1) Add 10ml DMSO to 90ml FBS.
2) Filter the mixture with a 0.2μm filter.
9. PBS (1×), calcium and magnesium free

### Isolation and culture of Primary Brown Preadipocytes and White Preadipocytes

1. Anesthetize the mouse using isoflurane.
2. Spark 70% ethanol on the mouse.
3. Using scissors to open the back midline from the neck.
4. Expose two lobes of the brown fat pad in the interscapular region.
5. Separate the interscapular brown adipose tissue (iBAT) and remove the surrounding tissues (Figure 1).
7. Dissect the subcutaneous white adipose tissue in the interscapular region (sWAT) and remove the surrounding tissues (Figure 1).
8. Keep the tissue in a 10 cm culture dish with ice-cold 1 X Phosphate Buffered Saline (PBS) on ice.
9. Transfer iBAT or sWAT into a 50ml falcon tube, place the tube horizontally, and place fat tissue near the open side.
10. Use scissors to mince fat tissue into small pieces (0.5-1 mm^3^).
11. Add 5-15ml digestion buffer (for iBAT or iWAT from one mouse) into a 50-ml falcon tube.
12. Place the tube in a horizontal position into a water bath with a shaker at 37 °C and 150 cycles/minutes. Digest iBAT for 45-60 min, sWAT for 30-40 min, vortex for 10 s every 15 min.
13. Terminate digestion by adding ¼ volume of FBS.
14. Pour the solution with digested tissue through a 100μm cell strainer into a new 50ml sterile tube.
15. Centrifuge at room temperature at 600g (or 1000 rpm) for 3-5 min.
16. Remove the supernatant.
17. Resuspend pellet in 3ml complete growth medium (for iBAT or sWAT from one mouse)
18. Plate the cells into one well of a 12-well plate, and keep the plates in a 37 °C, 5 % CO_2_ Incubator Day 0) overnight.
19. Next day (Day 1), wash cells with a pre-warmed growth medium for 2-3 times and add fresh primary preadipocytes growth medium. If the cells are intended to be in primary cell condition, change the growth medium every other day, if the cells are intended to be immortalized, perform the immortalization process immediately.

## IMMORTALIZATION OF MOUSE BROWN AND WHITE PREADIPOCYTES

On day 1 of preadipocyte isolation and primary culture, retroviral transduction needs to be performed to immortalize cells.

1. Prepare fresh culture media containing 8μg/mL polybrene.
2. Wash the cells with warmed culture medium for 2-3 times.
3. Add fresh culture media containing 8μg/mL polybrene (0.5ml/well for 12-well plate or 1ml/well for 6-well plate).
4. Add retroviral particle solution from the step of retroviral packaging, and incubate cells at 37°C, 5% CO_2._
To calculate the volume of virus needed to infect the target cells, it is necessary to obtain the parameters of target cell number, multiplicity of infection (MOI), viral titer, or viral infectious titer.
4.1 Multiplicity of infection (MOI) refers to the number of infecting viral particles per cell. MOI of each cell type or cell line varies. MOIs can be referenced from published literature or the test to determine the optimal MOI for infection.
4.2 Viral titer or viral particle concentration (VP/ml) obtained from retrovirus quantitation.
4.3 Functional titer or Infection titer refers to the number of viral particles capable of infecting targeted cells. It can be expressed as Transduction units per ml (TU/ml), infectious units per ml (ifu/ml, or plaque-forming units per ml (pfu/ml). The infection titer varies depending on the type of target cell lines or transduction methods (Kahl C.A. et al., 2004).
Viral particles needed for infection: targeted cell number × MOI
Viral particles volume needed for infection: viral particle needed / infectious titer
For example: the targeted cell number: 10^4^

MOI: 100
Infectious titer: 10^8^ ifu/ml, or 10^5^ ifu/μl
Therefore

Viral particle volume needed for infection: 10^4^ × 100 / 10^5^ = 10μl
Add 10μl packaged retrovirus to 1ml medium (well/6-well plate) or 0.5ml medium (well/12-well plate)
5. Viral infection to the cells for 24-48 hours.
6. Remove the virus-containing medium and replace it with a fresh culture medium.

Note: Polybrene increases the efficiency of viral infection. However, it is toxic to some cell lines. If the cell toxicity from polybrene is observed, substitute protamine sulfate for polybrene. If viral toxicity is observed in your cell line, decrease the infection time to between 4 to 20 hours.

**Figure 1.**
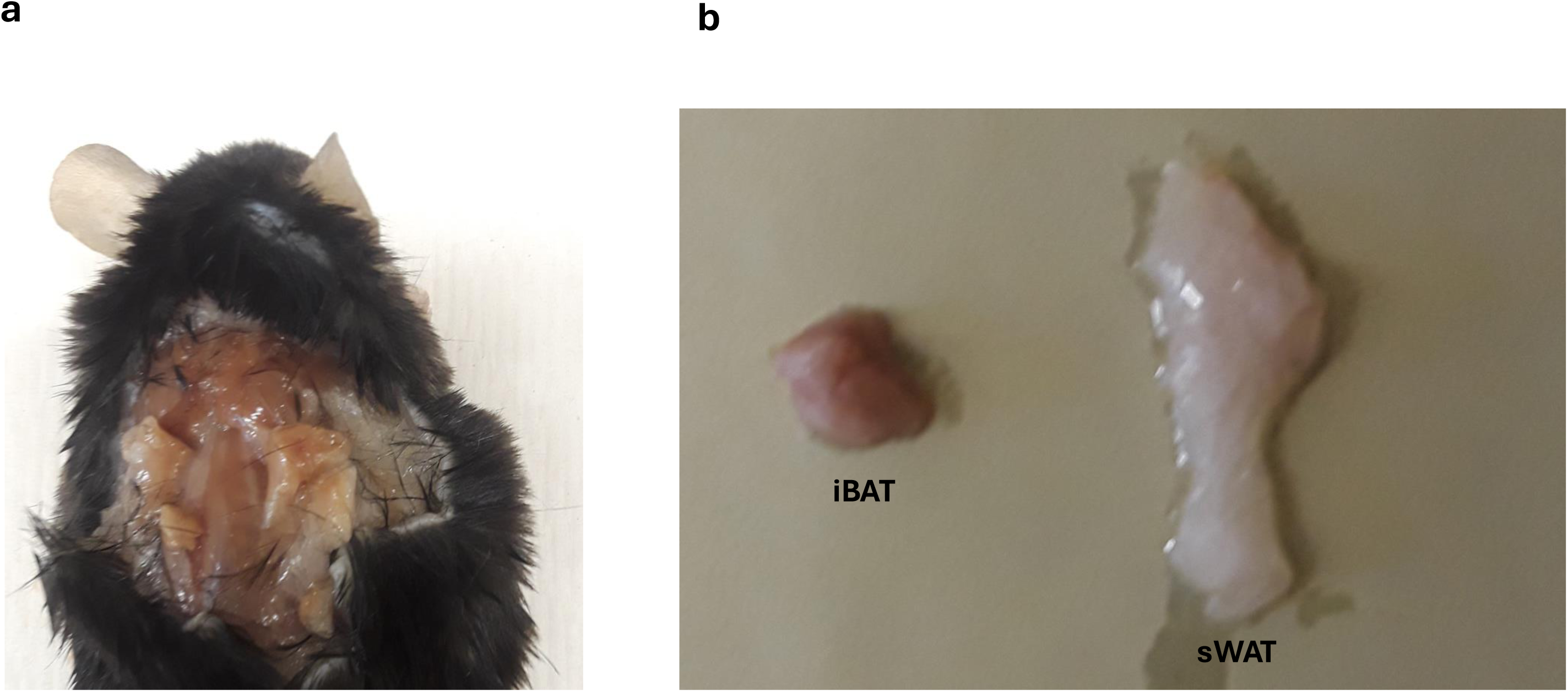
Mouse brown adipose tissue (BAT) and white adipose tissue (WAT). The brown adipose tissue was dissected from interscapular region (iBAT). The white adipose tissue was separated from subcutaneous white adipose tissue in the same region (sWAT). (a) depots of interscapular brown adipose tissue (iBAT), and subcutaneous white adipose tissue (sWAT); (b) separated interscapular brown adipose tissue (iBAT) and subcutaneous white adipose tissue (sWAT)

## SELECTION OF IMMORTALIZED MOUSE PREADIPOCYTES

1. After retroviral transduction, let preadipocytes grow in a normal growth medium at least for 48 hours, allowing resistant genes to express.
2. Wash cells with warmed 1× PBS for one time, add a growth medium containing antibiotic for selection only, and do not include any other antibiotic or antifungal reagents that may interfere with selection.
3. Replace antibiotic-containing medium for every other day, which also removes the dead cells. The selection process needs to continue for 7-10 days. Most non-resistant cells will be eliminated within 10 days, leaving the resistant cells to grow.
4. After the initial selection process, cells can maintain in normal growth medium without the selection antibiotic, which allows them to proliferate without selection pressure. However, to ensure the maintenance of stability of the integrated gene, cells can be occasionally, for example every 5 passages, sub-cultured in a medium containing selection antibiotic for a new round selection.
5. When growing to 70-80% confluence, the preadipocytes can be frozen as a polyclonal cell line or plated for selection of single-cell clones.

Note: the concentration of antibiotic to select and maintain can be referenced from published literature reports or perform kill curve assay to determine the optimal concentration.

## SELECTION OF SINGLE-CELL CLONES OF IMMORTALIZED MOUSE PREADIPOCYTES

1. Wash preadipocytes with 1× PBS and detach cells by trypsinization.
2. Dilute cells in an antibiotic-containing growth medium with cell concentration of 10 cells/ml.
3. Seed 100μl of cell suspension to each 96-well plate. Theoretically, each well will contain one cell.
4. Alternatively, after detaching the cells, sort a single cell to each well of the 96-well plate by cell cytometry.
5. Culture the cells overnight and mark the wells that contain a single cell.
6. Observe the cell growth in the marked wells that initially with a single cell.
7. When the cell grows and forms a single colony with a certain degree of confluence, transfer cells to a well of 12-well plate, let the cells continue to expand, then subculture cells to a well of 6-well plate.
8. Once the cells reach 70-80% confluence in a well of 6-well plate, cells can be transferred to multiple wells of the plate or a tissue culture flask for further expansion.
9. Freeze cells from the earliest passage as single-cell clones.

## CRYOPRESERVATION OF IMMORTALIZED MOUSE PREADIPOCYTES

1. Wash preadipocytes with 1× PBS for one time and add a couple of drops of trypsin to cover the cell surface.
2. Incubate the cells at 37 °C in the incubator for 2-3 minutes.
3. Add normal growth medium to the well (1ml / well, 6-well plate).
4. Detach and mix cells by pipetting up and down.
5. Collect cells to a new 50-ml tube.
6. Centrifuge the cells at 1000 rpm for 3 minutes.
7. Carefully remove the supernatant and avoid disturbing the cell pellet.
8. Add cell freezing medium to the tube and vortex for 30 seconds to resuspend cells, usually, 3ml freezing medium for one of 6-well plate cells.
9. Aliquot 1 ml of cells to one of 1 ml-cryotube.
10. Place cell cryotubes in a freezing container containing 100% isopropyl alcohol and keep them in a −80°C freezer overnight.
11. Storage the cells in liquid nitrogen for long-term reservation.

## THAWING AND CULTURE OF CRYOPRESERVED IMMORTALIZED MOUSE PREADIPOCYTES

1. Take cryotubes containing preadipocytes out of a liquid nitrogen tank.
2. Keep cryotubes in a 37°C water bath for 2-3 minutes to thaw the cells.
3. Centrifuge cryotubes at 1000 rpm for 3 minutes.
4. Discard supernatant (freezing medium) and do not disturb the cell pellet in the bottom of the tube, the cell pellet could be visible or invisible.
5. Add 1 ml of cell growth medium into each cryotube, and vortex for 10 seconds.
6. Transfer the 1 ml of cell suspension to a new 50 ml tube.
7. Add additional cell growth medium and vortex for 10 seconds.
8. Plate the cells in tissue culture plates, dishes, or flasks.
9. Culture the cells in growth medium at 37°C in a humidified incubator with 5% CO_2_.

## SUBCULTURE AND EXPANSION OF IMMORTALIZED MOUSE PREADIPOCYTES

When the preadipocytes reach 80-90% confluence, subculture cells with a ratio of 1:3 to 1:4.

1. Aspirate culture medium, wash cells with 1× PBS for one time.
2. Add 0.25% (w/v) trypsin-EDTA to cover the cell surface.
3. Incubate the cells in an incubator at 37°C for 2-3 minutes.
4. Observe cells under an inverted microscopy with 4× or 10× magnification. When fibroblast-shaped preadipocytes shrink and become dot-shaped cells, add growth medium to inactive trypsin (1ml / well of a 6-well plate).
5. Detach cells by pipetting up and down, and then collect cells to a new 50-ml tube.
6. Centrifuge the cell suspension at 1000 rpm for 3 minutes.
7. Carefully discard the supernatant (trypsin and medium), do not disturb the cell pellet in the bottom of the tube. The cell pellet may be visible or invisible.
8. Add growth medium to the tube, and vortex for 10 seconds, usually one plate of 6-well preadipocytes can split into 3-4 plates of 6-well plates, add growth medium accordingly.
9. Plate the cells in new tissue culture plates, dishes, or flasks.
10. Culture the cells in the incubator at 37°C, with 5% CO_2_, to allow the cells to grow.
11. Replace with fresh growth medium every other day. When the cells reach 80-90% confluence, either subculture cells again or freeze cells for cryopreservation.

## DIFFERENTIATION OF IMMORTALIZED PREADIPOCYTES

1. Grow the preadipocytes in the growth medium, change the medium every other day.
2. When the cells reach 100% confluence, change the medium with the freshly prepared induction medium as differentiation day 0.
3. Keep the cells in an induction medium for 2-3 days, then change the medium with a freshly prepared differentiation (or maintenance) medium, replacing the differentiation medium every other day until differentiation day 12.
4. After preadipocytes differentiate into mature adipocytes, the cells can be used for designed experiments such as treatment of testing drugs, reagents, etc., or harvest or freeze differentiated adipocytes for further assays.

Note: The lipid droplets will appear after induction and will significantly accumulate during the differentiation period. UCP-1 protein, one of the brown adipocyte markers, can be detected in 4-5 days of post-induction.

## IDENTIFICATION OF DIFFERENTIATED WHITE ADIPOCYTES AND BROWN ADIPOCYTES

### CELL MORPHOLOGY

Preadipocytes are the fibroblast-like shape and will turn round-shaped cells during and after induction. Multiple lipid droplets with different sizes can be observed under microscopy after induction and significantly accumulate during differentiation period (Figure 2, Figure 3).

**Figure 2.**
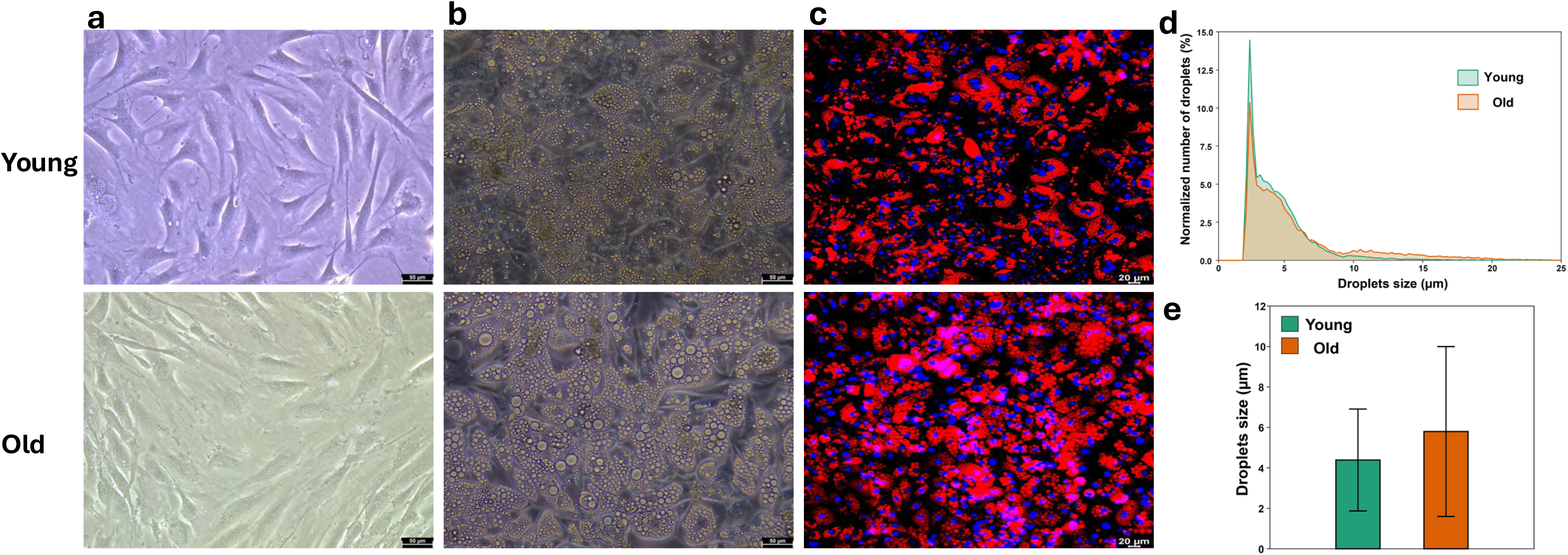
Morphology of immortalized brown preadipocyte and differentiated brown adipocyte. The stromal vascular fraction (SVF) cells were isolated from interscapular brown adipose tissue (iBAT) of young (1-day-old pups) and old (27-month-old) mice. Cell images were acquired using Leica DMi8 microscopy with Leica Application Suite X software. The magnification was 10 x 20. (a) cells in differentiation day-0 under visible light; (b) cells in differentiation day-12 under visible light; (c) fluorescence image of lipid droplets and nuclei in differentiation day-12, staining with lipid TOX deep red neutral lipid stain and Hoechst 33342 respectively. (d) Quantification of lipid droplet size distribution and (e) average droplet size of old and young differentiated brown adipocytes show distinct differences between cells isolated from mice of different ages.

**Figure 3.**
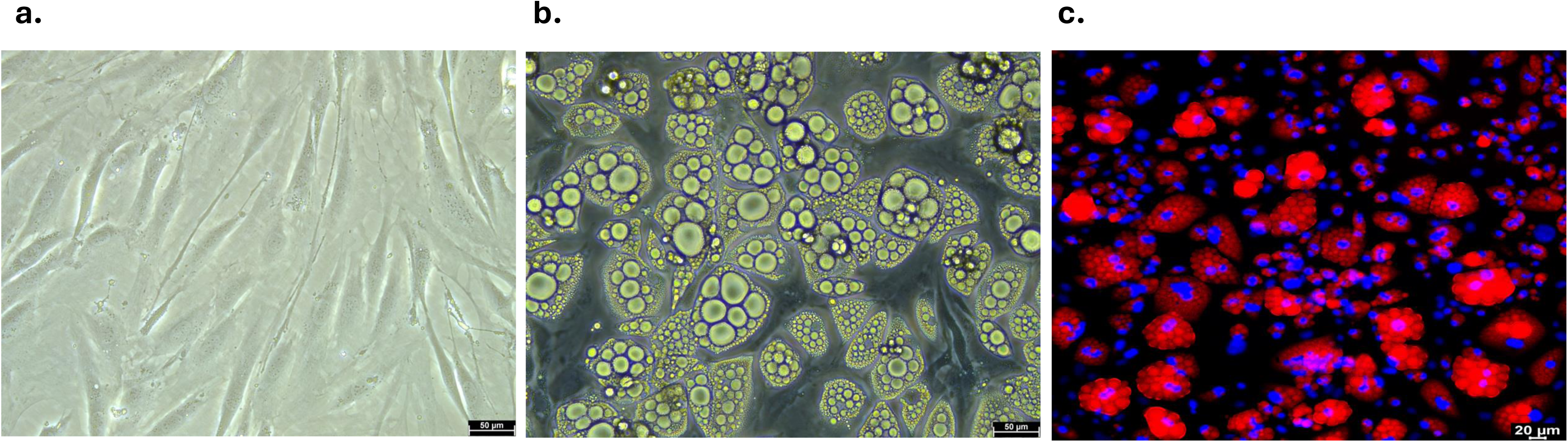
Morphology of immortalized white preadipocyte and differentiated white adipocyte. The stromal vascular fraction (SVF) cells were isolated from mouse subcutaneous white adipose tissue in the interscapular region (sWAT) of 12-month-old mice fed with high fat and high fructose diet for 10 months. Cell images were acquired using Leica DMi8 microscopy with Leica Application Suite X software. The magnification was 10 x 20. (a) cells in differentiation day-0 under visible light; (b) cells in differentiation day-12 under visible light; (c) fluorescence image of lipid droplets and nuclei in differentiation day-12, staining with lipid TOX deep red neutral lipid stain and Hoechst 33342 respectively.

### LIPID DROPLET STAINING AND NUCLEUS COUNTERSTAINING

Lipid droplets in live differentiated adipocytes can be stained by lipid-specific dyes, for example LipidTOX^TM^ neutral lipid stains, for monitoring adipogenesis. Cell nuclei DNA can be stained by Hoechst 33342 (Figure 2, Figure 3).

#### Materials

HCS lipidTOX^TM^ Deep Red neutral lipid stain, Invitrogen/Thermo Fisher Scientific #: H34477

Hoechst 33342, 20mM in stock, Thermo Scientific #: 62249

1. Prepare Hoechst working solution (1mg/ml). Dilute 9μl of Hoechst 33342 (20mM stock solution) into 91μl 1×PBS as a working solution (1mg/ml).
2. Prepare lipid and nuclei staining medium. Add 5μl of HCS lipidTOX^TM^ Deep Red neutral lipid stain and 1μl of Hoechst working solution to 1ml cell culture medium, and mix staining medium well. Scale up the volume as each experiment needs

Note: HCS lipidTOX^TM^ Deep Red neutral lipid stain is diluted with 1:200 in staining medium. Hoechst nuclei stain has a final concentration of 1μg/ml in the staining medium. The final staining concentration for both lipid staining and nuclei staining needs to be optimized for each experiment.

3. Aspire cell culture medium, add stain medium (1ml per well for six-well plate).
4. Incubate cells in a cell culture incubator for 30 - 40 minutes.
5. Immediately monitor and image the lipid droplet staining using fluorescence microscopy with appropriate filters.

### LIPID DROPLETS ANALYSIS

Lipid droplet number, lipid size, and distribution can be analyzed using open-source image J software based on cell microscopy images.

Alternatively, Lipid droplet number, lipid size, lipid diameter, and lipid mass quantification can be achieved by optical diffraction tomography and Raman spectroscopy (Anantha et al., 2024)

### DETECTION OF ADIPOGENESIS MARKERS

Using qPCR and/or Western Blot to detect adipogenesis markers such as PPARγ, FABP4 (aP2), etc. These proteins should be expressed in both differentiated brown adipocytes and white adipocytes (Figure 4).

**Figure 4.**
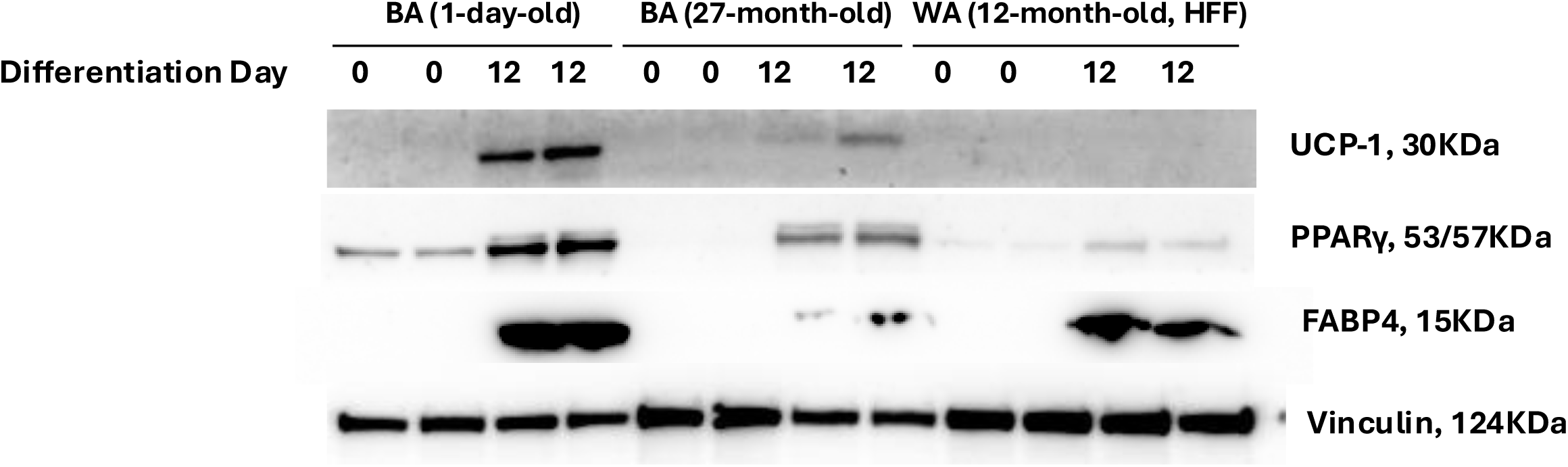
Expression of UCP-1, PPARγ, FABP4 protein by western blot in differentiated brown adipocytes (BA) and white adipocytes (WA) at day-0 and day-12. Brown preadipoctyes were isolated from 1-day-old pups or 27-month-old mice. White preadipocytes were isolated from 12-month-old mice fed with high fat and high fructose diet (HFF) for 10 months. Vinculin served as house keeping protein.

### DETECTION OF BROWN ADIPOCYTE MARKERS

Using qPCR and/or Western Blot to detect brown adipocyte markers. UCP-1 is the standard marker for brown adipocytes (Figure 4). PGC-1α or PRDM16 are also considered brown adipocyte markers.

### MITOCHONDRIAL STAINING AND NUCLEUS COUNTERSTAINING

Mitochondria can be stained by mitochondria specific dyes, for example, MitoTracker deep red FM for monitoring mitochondria biogenesis. Cell nuclei DNA can be stained by Hoechst 33342 (Figure 5, Figure 6).

**Figure 5.**
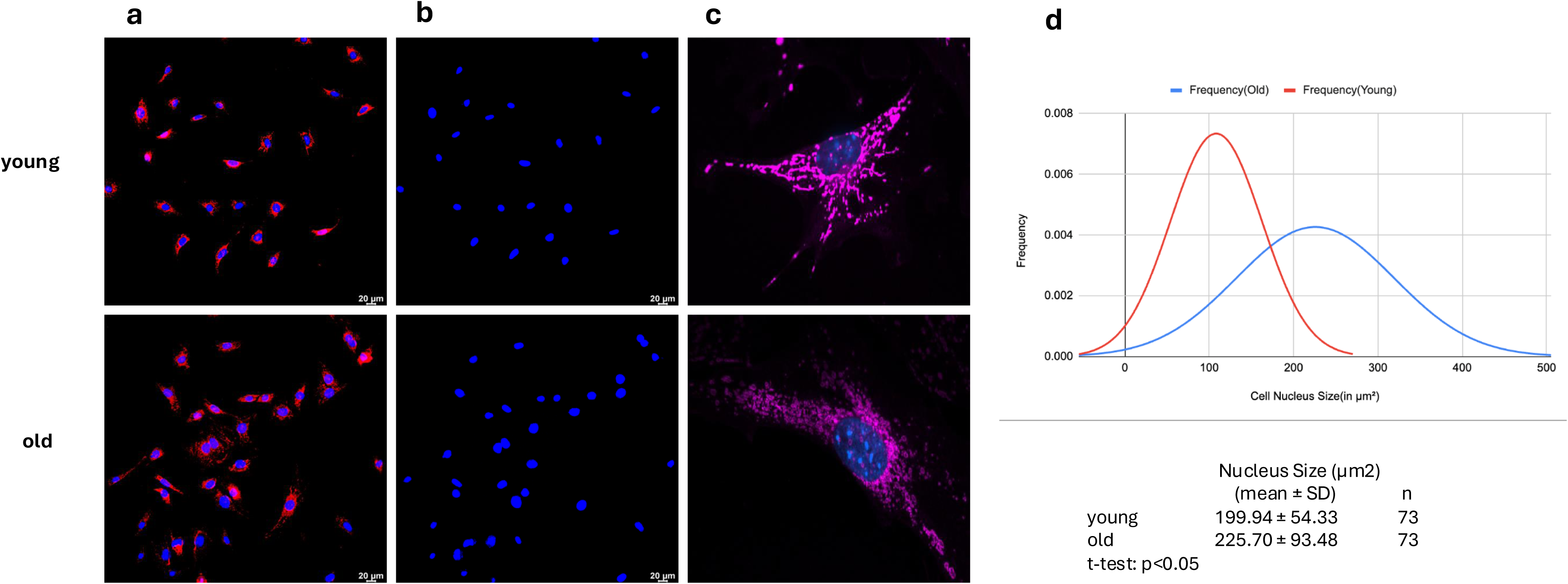
Mitochondria morphology and nucleus size of young and old immortalized brown preadipocytes. The brown preadipocytes were isolated from iBAT of young (1-day old pup) and old (27 months) mice. (a) fluorescence images of mitochondria and nuclei and (b) fluorescence images of nuclei only were acquired using Leica DMi8 microscopy with magnification of 10 x 20. (c) representative images of mitochondria and nucleus of individual cells, which were acquired using confocal microscopy of Nikon W-1 Spinning Disk with magnification of 10 x 60; (d) distribution of cell nucleus size of young and old preadipocytes.

**Figure 6.**
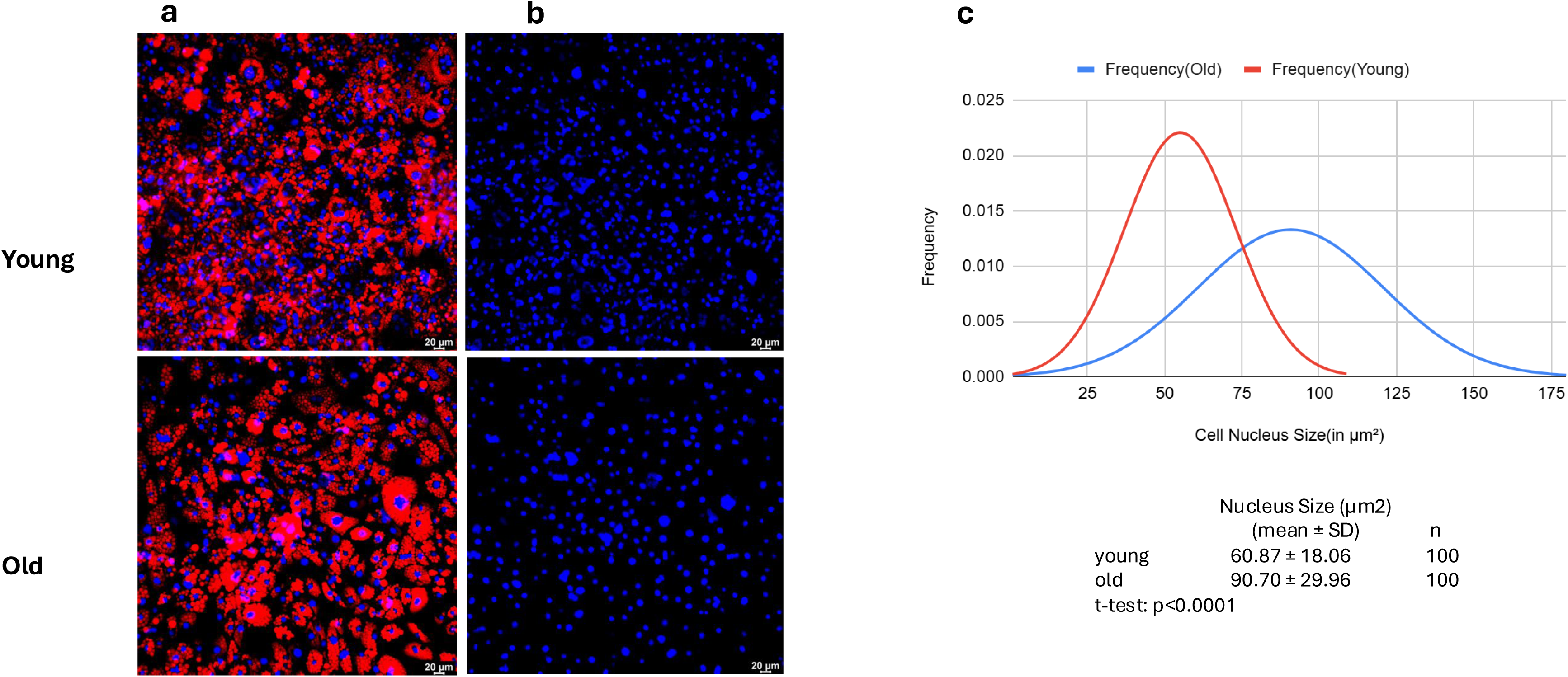
Mitochondria morphology and nucleus size of young (1-day-old) and old (27-month-old) differentiated brown adipocytes. (a) fluorescence images of mitochondria and nuclei and (b) fluorescence images of nuclei only were acquired using Leica DMi8 microscopy with magnification of 10 x 20; (c) distribution of cell nucleus size of young and old differentiated adipocytes.

#### Materials

MitoTracker deep red FM, Cell Signaling # 8778, 50ug/vial

Anhydrous dimethyl sulfoxide (DMSO)

Hoechst 33342, 20mM in stock, Thermo Scientific #: 62249

Note: other MitoTracker probes such as MitoTracker green FM, MitoTracker orange can be applied as well. Freshly prepare the staining solution just before use.

1. Prepare 1mM stock solution of MitoTracker deep red FM Briefly centrifuge the MitoTracker vial, add 91.98μl DMSO to a vial containing 50μg MitoTracker deep red FM, vortex the vial, and make sure MitoTracker is completely dissolved in DMSO.
2. Prepare Hoechst working solution (1mg/ml) Dilute 9μl of Hoechst 33342 (20mM stock solution) into 91μl 1×PBS as a working solution (1mg/ml).
3. Prepare staining solution Add 0.5μl MitoTracker stock solution and 1μl Hoechst working solution to 1ml culture medium without serum. The final concentrations of MitoTracker and Hoechst are 500nM and 1μg/ml respectively.

Note: Adjust the volume according to each experiment’s needs. The final staining concentration for both MitoTracker and Hoechst nuclei staining varies for different cell types and different conditions. The staining concentration needs to be optimized for each experiment.

4. Staining the cells Cells are cultured on cell culture plates, dishes, or coverslips. Remove media, add prewarmed staining solution, and incubate cells (37°C, 5% CO_2_) for 30 min. After incubation, remove staining media, add fresh prewarmed media or buffer, and observe and image cells alive using fluorescent microscopy immediately, or fix cells first and then observe and image the cells.

Note: incubation times depend on the cell types or experiment conditions. Usually, 15-45 minutes are sufficient. The incubation times need to be optimized for each experiment.

5. Fixation after staining (Optional) After staining, remove staining media, fix cells in ice-cold 100% methanol for 15 min at - 20°C, rinse cells 3 times with 1X PBS, and observe and image cells using fluorescent microscopy.

### CELL AGING EVALUATION

Several aging markers have been established in in vitro cell culture (Hartmann et al., 2023), which can generate either individual values or patterns to estimate donor’s age. Here we demonstrated that aged cells exhibit enlarged nuclei size and fragmented mitochondria (Figure 5, Figure 6).

## COMMENTARY

### Background Information

#### Immortalized cell lines

Primary cells are more closely represented in vivo tissues because they are directly isolated from tissues. Primary cells also keep genetic integrity, providing a valuable and high physiological relevance (Husna et al., 2022; Severgnini et al., 2012). However, after some generations, primary cells will lose their original biologic features and reach senescence. Repeated generation of fresh primary cells is time-consuming, high cost, and can cause significant variation from donor to donor. Immortalized cells can repeal normal cellular senescence and maintain cell division due to natural mutation or intentional genetic modification. Immortalized cells can retain the phenotype in their parent tissues, which makes them a powerful tool for studying cell growth, metabolism, and senescence. It also needs to acknowledge that multiple factors can influence the extent of phenotype retention in immortalized cell lines including immortalization methods, the specific cell type, and the genetic stability of the resulting cell lines (Tani et al., 2024; Katwal et al., 2019). Several strategies are practical to make cells immortal. (1) spontaneously immortalized cell lines. Primary cells in some conditions can naturally undergo genetic mutation to confer a growth advantage, which allows the cells to invalidate normal cellular senescence and apoptosis. The best example of a spontaneous immortalized cell line is 3T3-L1 cells commonly used for studying adipogenesis. (2) expression of viral tumor suppressor genes that revoke the cell cycle and overcome senescence. The most applied of these genes are the Large T-antigen of the Simian virus (SV-40) and human papilloma virus (HPV) E6 and E7. They both suppress retinoblastoma (Rb) and P53 genes which are both crucial for cell cycle controlling. The well-known and widely used HEK293T cell line is immortalized by the expression of SV40 large T-antigen. (3) Using gene editing technology such as Crispr/Cas9 to directly inactivate specific tumor suppressor genes (e.g. retinoblastoma (Rb) and P53) to promote cell immortality. (4) Expression of human telomerase reverse transcriptase (hTERT) to upregulate telomerase. The ribonucleoprotein telomerase can preserve telomere length which otherwise will be shortened with each cell division. Thus, cells can escape replicative senescence and continue dividing indefinitely. Here we express SV40 large T-antigen delivered by retrovirus to immortalize brown and white preadipocytes.

#### The components of the induction medium and maintenance medium

Inducing differentiation of brown adipocytes from precursor cells such as brown preadipocytes isolating from brown adipose tissue in vitro requires a specific induction medium that contains some key components to promote differentiation and maturation. Besides basic medium DMEM/F12 and FBS, the typical induction medium for brown adipocytes includes six components. (1) Insulin: stimulates glucose uptake and promotes the expression of genes involved in adipogenesis and brown adipocyte differentiation; (2) Dexamethasone: A glucocorticoid that promotes adipogenic differentiation by activating certain transcription factors such as CCAAT/enhancer-binding protein (C/EBP); (3) Indomethacin: A nonsteroidal anti-inflammatory drug (NSAID) that can promote brown adipocyte differentiation; (4) 3-Isobutyl-1-methylxanthine (IBMX): A phosphodiesterase inhibitor that increases cyclic AMP (cAMP) levels, leading to the activation of protein kinase A (PKA) and subsequent activation of adipogenic and thermogenic transcription factors; (5) Rosiglitazone: promotes the differentiation of preadipocyte into mature adipocytes by activating peroxisome proliferator-activated receptor gamma (PPARγ); (6) Triiodothyronine (T3): A thyroid hormone that is critical for brown adipocyte differentiation and the activation of thermogenic genes, such as uncoupling protein 1 (UCP1). White adipocyte induction medium includes everything else with brown adipocytes except Triiodothyronine (T3), while indomethacin is optional. Biotin and pantothenate can be optionally included in the induction medium since both are necessary for fatty acid synthesis and overall cell metabolism. The maintenance medium followed by induction includes a growth medium supplement with insulin and T3 for brown adipocytes, and insulin for white adipocytes, which continue to support cell survival, functions, and differentiation (Shamsi and Tseng 2017; Cero et al., 2021; Wu et al., 2005; Boucher et al., 2012).

#### White adipocytes vs. brown adipocytes

White adipocytes typically have large cell sizes, harbor a single large lipid droplet, and have fewer mitochondria. White adipocytes function as energy storage and adipokines secretion (such as leptin, adiponectin, TNFα, IL-6, etc.) to regulate appetite, energy metabolism, and immune response, and have lower metabolic activity (Morigny et al., 2021). In contrast, brown adipocytes retain small cell size and contain multiple small lipid droplets and abundant mitochondria. Brown adipocytes serve as energy expenditure, maintaining body temperature through non-shivering thermogenesis that is mediated by uncoupling protein 1 (UCP-1) in mitochondria. Brown adipocytes have a higher metabolic rate and also secrete adipokines (such as adiponectin, IL-6, norepinephrine, VEGF, etc.) to regulate energy metabolism, immune response, and thermogenesis (Svensson et al., 2011; Li et al., 2021; Wang et al., 2014; Cypess, 2022). We particularly studied the nuclei size of brown adipocytes. Our data (Figure 5, Figure 6) indicates nuclei size of brown preadipocytes is larger than that of differentiated brown adipocytes, suggesting brown adipocyte nuclei compacted after differentiation.

#### Young adipocytes vs old adipocytes

Young adipocytes preserve smaller cell size, accumulate less lipid, maintain favorable integrity of the cellular structure, efficiently manage lipid storage and mobilization, secrete beneficial adipokines such as adiponectin, and own better vascularization. All of these contribute to maintaining overall healthy metabolic homeostasis. Yet, old adipocytes possess larger cell sizes, undergo more dysfunctional lipid metabolism and adipokine secretions, and escalate cellular senescence, which is attributed to slow metabolic rate, inflammation, insulin resistance, and metabolic disorders. The structure and functional alterations of old adipocytes significantly deteriorate the aging process and aging-related metabolic diseases (Muller et al., 2011; Cartwright et al., 2010; Wu et al., 2013; Lumeng et al., 2011).

Several biological indicators could serve as aging markers, particularly in the in vitro cell culture system (Hartmann et al., 2023). For example (1) primary aging markers: DNA damage, histone modification, telomere attrition; (2) cellular senescence markers: senescence-associated-β-galactosidase (SA-β-GAL), senescence-associated secretory phenotype (SASP) such as IL-6, IL-8, MCP-1, etc.; (3) antagonistic aging markers: Lamin B1 expression, nucleus size, etc. Here we use nucleus size and mitochondria structure integrity to identify the young or old preadipocytes or differentiated adipocytes. We are undergoing senescence-associated secretory phenotype (SASP) and RNA sequencing to further characterize the young and aged adipocytes.

#### Staining mitochondria, nuclei, and lipid droplets in live adipocytes vs. fixed cells

Choosing non-toxic or low-toxicity dyes for live-cell staining enables real-time observation of the dynamics and functions of mitochondria, nuclei, and lipid droplets without compromising their integrity or introducing artifacts from fixation. Fixatives can distort the structure and size of mitochondria (Qin et al., 2021; Kim et al., 2022; Richter et al., 2018; Zhang et al., 2017) and nuclei (Zarebski et al., 2021; Solovei et al., 2002; Deal & Henikoff 2010), as well as alter the size and distribution of lipid droplets due to lipid extraction or deformation (Xu et al., 2017; Christianson et al., 2010; Lumaquin et al., 2020; Huang et al., 2023). However, if high-resolution imaging and detailed structural analysis are the primary goals, staining in fixed cells is the preferred option.

### Critical Parameters

#### Retrovirus production

To effectively package the viral vectors, we seeded 293T cells around 95%, or around 1.2 × 10^6^ cells/well for a 6-well plate followed by transfection the next morning. The first collection of viral containing medium is done at 24 hours of post transfection, and the second collection is done at 56 hours of post-transfection. To maximize the virus harvest, a viral-containing medium also can be collected 3 times every 24 hours.

#### Isolation of primary brown and white preadipocyte

Collagenase type II can effectively degrade the collagen in adipose tissues and dissociate tissue into separated individual cells in suspension with minimized cell damage (Sharun et al., 2021; Pak et al., 2016). It is crucial to choose collagenase type II for adipose tissue digestion. We found brown adipose tissue needs more digestion time than white adipose tissue, and old adipose tissues are more difficult to digest than young adipose tissue. Brown adipose tissues from one-day-old pups are very easy to digest. We digested brown adipose tissue for 45-60 minutes and white adipose tissue for 30-40 minutes. The centrifuge speed for pelleting suspended cells is 600 g or 1000 rpm to minimize cell damage.

#### Culture of primary brown and white preadipocyte

The components of the cell culture medium could be varied depending on the purpose of the experiments. For cellular energy metabolism, attention forces on the glucose concentration in the medium. High glucose (usually 25mM) medium can be applied to cancer cell lines such as HepG2 cells, and certain immortalized cell lines such as HEK293 cells because these cells have high metabolic rates and high energy demands. High glucose can support rapid cell growth and high proliferation rates (Iyer et al., 2010; Furuichi et al., 2021; Iuchi et al., 2010). However, high glucose can alter cellular metabolism and cause cell stress responses (Cong et al., 2020; Gerlini et al., 2018; Palacios-Ortega et al., 2015; Krycer 2020; Chen 2017). This consideration should be taken when interpreting experiment results. Low glucose (usually 5mM) medium is more suitable for primary cells, mesenchymal stem cells, and certain types of differentiated cells such as adipocytes because these cells naturally exist in lower glucose environments. Low glucose reduces the risk of metabolic alterations and stress responses and closely mimics physiological levels found in most tissues (Liu et al., 2020; Pham & Pham, 2019; Wang et al., 2016; Lakhkar et al., 2015; Hirasawa et al., 2014; Wang et al., 2015). We use nutrient-enriched DMEM/F12 with intermediate glucose level (3.15g/L) supplemented with 10% FBS and antibiotics as primary brown and white preadipocyte growth medium, which is balanced on nutrient support for cell attachment and growth and the impact of high glucose on cellular metabolism and stress response.

#### Immortalization of brown and white preadipocyte

Polybrene can reduce the charge repulsion between the viral particles and the cell surface to increase the efficiency of retrovirus transduction (Landazuri et al., 2007; Han et al., 2015). However, some cell types are sensitive to polybrene which may cause cell toxicity and cell death. The concentration of polybrene needs to be optimized for each experiment. By using 8μg/ml of polybrene, we did not observe any cell toxicity and cell death for brown and white preadipocytes. To efficiently infect cells, the volume (or number) of viral particles to infect targeted cells must be calculated based on the targeted cell number, virus infectious titer, and multiplicity of infection (MOI) which must be tested and optimized for each cell type and each experiment. The MOI information also could be referenced from the literature. The MOI value we used is 100, which is for the retrovirus we packaged to infect mouse brown and white preadipocytes that we isolated and cultured. The viral transduction is highly efficient with this MOI value.

#### Differentiation of brown and white preadipocyte

The immortalized primary brown and white preadipocyte first needs to be induced to differentiate for adipocytes. The concentrations of dexamethasone, indomethacin, IBMX, rosiglitazone and T3 in the induction medium are quite consistent across the protocols, however, insulin concentrations vary from 18nM to 1720nM, (Singh et al., 2015; Wang et al., 2018; Choi et al., 2013; Park et al., 2013; Wu et al., 2005; Cero et al., 2021; Shamsi and Tseng et al., 2017; Boucher et al., 2012) in the literature. Based on our and other experiments and results, 20nM insulin is sufficient in the induction medium for inducing both brown and white preadipocytes into differentiation (Boucher et al., 2012; Choi et al., 2013; Elsaid et al., 2024). Optimal insulin concentration is essential for effective differentiation. More insulin is not necessarily better.

Once insulin receptor is saturated, excessive insulin will bind to insulin-like growth factor 1 receptor (Zhou et al., 2018; Menting et al., 2015) and hybrid receptor (Pandini et al., 2002; Slaaby et al., 2006) or interact with other receptors such as insulin receptor-related receptor (Deyev et al., 2011), G-protein coupled receptors (Hyun et al., 2010; Meister et al., 2014; Fu et al., 2014), or scavenger receptor class B type 1 (Tondu et al., 2005; Cavallari et al., 2018). These will cause cellular stress, stimulate cell proliferation, alter the metabolic response, and undermine experimental reproducibility and consistency.

#### Centrifuge speed for live cell pelleting

The centrifuge speed for pelleting live cells in any step of cell culture handling must be around 500 – 600 g (or 800 – 1000 rpm) in a short period (3 minutes). High-speed centrifuge can cause cell membrane deformation or rupture, and deceleration after high-speed centrifuge can generate shear force that leads to cell lysis or mechanical damage.

### Troubleshooting

Table 1 – 3 describes the problems encountered in conducting basic protocols 1 – 5 and their possible causes and solutions.

**Table 1.** Potential problems, causes and troubleshooting in establishing immortalized mouse preadipocyte cell lines.

| Problem | Possible cause | Solution |
| --- | --- | --- |
| Low titer of retrovirus that generated | HEK293T cells in culture for long period time with high passages or overconfluent cells. | Using freshly thawed HEK293 cells with low passages, never overgrow and overconfluent cells for subculture and transfection. |
|  | HEK293T cells in insufficient culture medium or incorrect temperature or CO<sub>2</sub> concentration. | Keep culture medium in a relative high level, for example at least add 3 ml medium / well for a 6-well plate, and frequently (1-2 days) change the medium, Check or calibrate the incubator in correct working status. |
|  | Poor-quality transfection reagent. | Do not leave transfection reagent in room temperature for too long time, make sure the reagent does not be contaminated with another reagent. |
|  | Inappropriate ratio of transfection reagent and plasmid DNA. | Check the reagent and plasmid DNA in the suggested ratio or run a test to optimize the ratio. |
|  | Suboptimal transfection condition. | Check culture medium color, incubator temperature and |
|  |  | CO2 level to make sure everything is in suggested condition. |
|  | Low-quality plasmid DNA. | Use high quality DNA preparation methods, ensure DNA is pure and in high concentration. |
|  | Inadequate harvesting timing. | Do not harvesting viral particles too early or too late. |

Table 1 (continue)
| Problem | Possible cause | Solution |
| --- | --- | --- |
| Low attached cell number after culture of isolated cells | Inappropriate digestion enzyme. | Make sure to use collagenase type II to digest adipose tissue. |
|  | Inappropriate digestion temperature, and shaking speed, insufficient digestion time. | Check the temperature and speed of water bath shaker, make sure it run at 37C and 150 cycles/minutes. Make sure to digest tissue for suggested time, if the cell number is still low, prolong the digestion time, but no longer than 2 hours. |
| No attached cells after culture of isolated cells | Wrong cell culture medium or culture conditions. | Check the culture medium that should be DMEM or DMEM/F12 with 10% or more FBS. Check and calibrate the cell culture incubator, make sure it runs in right conditions. |
| Cell got contaminated | Insufficient wash initial attached cells culture medium that including many dead cells, blood cells or potential bacteria, fungi contamination. | Remove initial culture medium next day, wash attached cells for multiple times to eliminate all dead cells, blood cells or potential bacteria, fungi contamination. |
|  | Use contaminated culture medium, or do not include | Use fresh opened medium, filter the medium after mix |
|  | antibiotic in the medium. | DMEN/F12 with FBS and antibiotics, make sure to include antibiotic primocin for primary culture. |
|  | Cell contaminated during process of culture. | Review every step and working environment of cell culture, make sure to follow the requirements of aseptic techniques. |

Table 1 (continue)
| Problem | Possible cause | Solution |
| --- | --- | --- |
| Too many cells or all cells are dead after antibiotic selection | Low efficiency of retrovirus transduction. | Check polybrene concentration in virus infection medium, make sure it is in optimal range. |
|  | Insufficient viral particles for infecting target cells. | Check the viral infection titer, multiplicity of infection (MOI) and targeted cell number, make sure add enough viral particles to infect target cells. |
|  | Add antibiotics for selection too early. | At least let the cells grow in normal growth medium for 48 hours of post transduction to allow resistant gene to express. |
|  | Antibiotics concentration is too high for selection. | Run a kill curve test to determine the optimal antibiotics concentration for selection. |

### Time Consideration

#### Support protocol 1

The timeframe for retrovirus production and quantification is about 4-5 days.

- Preparation of 293T cells: overnight or 18 hours
- Plasmid transfection: 8 hours
- Collection of viral production: 24-72 hours
- Retrovirus quantification: 3 hours

#### Basic protocol 1

- Prepare the reagents, solutions and media: 4-6 hours
- Isolation of primary brown and white adipocyte: 3 hours
- Primary cell culture: overnight, or 18 hours

#### Basic protocol 2

- Retrovirus transduction and infection: 48 hours

#### Basic protocol 3

- Selection of immortalized preadipocytes with antibiotics: 10 days

#### Basic protocol 4

- Selection of single cell clone of immortalized preadipocytes: 3-5 weeks

#### Support protocol 2

- Cryopreservation of immortalized preadipocytes: 1 hour

#### Support protocol 3

- Thawing and culturing cryopreserved cells: 30 – 60 minutes

#### Support protocol 4

- Subculture immortalized preadipocytes: 1-2 hours
- Expansion immortalized preadipocytes: 3 – 6 days

#### Basic protocol 5

- Differentiation of brown preadipocytes to mature brown adipocytes: 12 days
- Differentiation of white preadipocytes to mature white adipocytes: 8 – 10 days

#### Support protocol 5

- Lipid staining and nuclear counterstaining: 1 hour
- Cell image: 1 – 3 hours

#### Support protocol 6

- Mitochondria staining and nuclear counterstaining: 1 hour
- Cell image: 1 – 3 hours

## Acknowledgments

We sincerely appreciate Dr. Christopher W. Ward and Dr. Jeanine A. Ursitti from the Department of Orthopedics, University of Maryland School of Medicine for providing brown and white adipose tissues from young (6-month-old) and aged (27-month-old) mice. We sincerely appreciate Confocal Microcopy Core (CMC) at the University of Maryland School of Medicine for technical support and assistance on cell imaging.

## Author Contributions

**Xiangdong Wu:** Methodology; Investigation; Data curation; Formal analysis; Writing – original draft. **Salaheldeen Elsaid**: Investigation. **Florian Levet:** Formal analysis **Winson Li**: Investigation. **Sui Seng Tee**: Methodology; Investigation; Data curation; Funding acquisition.

## Data Availability Statement

All data is included in this protocol.

